# Differential AXL expression and regulation of Arf1 controls matrix stiffness-dependent Golgi organization and function in breast cancer cells

**DOI:** 10.1101/2025.02.21.639478

**Authors:** Arnav Saha, Tushar Sherkhane, Nagaraj Balasubramanian

## Abstract

Integrin-mediated cell-matrix adhesion regulates cell growth and survival and is often deregulated in transformed cancer cells, promoting anchorage independence. Integrins are essential regulators of cellular mechanotransduction, known to be altered in cancer cells. In breast cancers, tumour stiffening is a well-characterized feature of the disease progression. The Golgi apparatus, regulated by integrin-mediated adhesion in responding to mechanosensory cues, could support cancer progression. MDAMB231 and MCF7 cells with distinct Golgi organization and altered mechanosensing were evaluated. In MDAMB231 cells the organized Golgi in stable adherent cells becomes disorganised on loss of adhesion. This contrasts with MCF7 cells, where the Golgi remains disorganized regardless of the adhesion status. In MDAMB231 cells, increasing matrix stiffness promotes Golgi organization and regulates Golgi-dependent microtubule acetylation. In MCF7 cells, the Golgi stays disorganised despite increasing stiffness. Both cell types show stiffness-dependent cell spreading, with MCF7 cells spreading more efficiently than MDAMB231 cells - a difference that may be partially mediated by their differential Golgi organization. AXL, a receptor tyrosine kinase, is known to be involved in rigidity sensing and is differentially expressed in MDAMB231 (high) vs MCF7 (no expression) cells. AXL inhibition (by R428) and siRNA-mediated AXL knockdown disrupt stiffness-dependent Golgi organization in MDAMB231 cells, promoting cell spreading. Transient and stable AXL expression in MCF7 cells causes the Golgi to become predominantly organised and respond to higher matrix stiffness, affecting cell spreading. Stiffness-dependent AXL expression supports stiffness-dependent Arf1 expression and activation to drive Golgi organization. Thus, matrix mechanosensing through the AXL-Arf1-Golgi pathway could regulate vital cellular processes in breast cancer cells.

## INTRODUCTION

Breast cancer is the most common invasive cancer in women, affecting approximately 12% of women worldwide (1). Physical cues like an increase in matrix stiffness, solid stress or an increase in the interstitial fluid pressure in the tumour microenvironment are implicated in breast tumour progression (2). A key physical hallmark of breast cancer progression is matrix stiffening, which contributes extensively to malignant transformation (3). Average breast tissue stiffness ranges from 0.5 kPa to 2.5 kPa, whereas breast tumor tissue stiffness could vary from 4 kPa to beyond 20 kPa (4).

Cells within the physiological niche have the inherent ability to sense and respond to the mechanical characteristics of the ECM, which govern an array of cellular processes, including migration, proliferation, differentiation, and spreading (5). Mechanical cues are also relayed inside the cells and thus could impact many pathways, including the dynamic organization of distinct intracellular compartments like the endomembrane organelles (6). Sensing of mechanical cues by cells is primarily done by Integrins (7), which regulates cellular mechanotransduction pathways (8). Integrin-mediated cell-ECM adhesion regulates crucial cellular processes like survival, growth, and proliferation, often deregulated in diseases like cancer (9). Previous work from our lab has shown that integrin-mediated cell-matrix adhesion is vital in regulating Golgi organization and functions in normal anchorage-dependent cells (10). Upon loss of matrix adhesion and loss of integrin signaling, Golgi undergoes dramatic disorganization in anchorage-dependent mouse fibroblasts. This is mediated by adhesion-dependent regulation of Arf1 activation (10).

Golgi organization and function are reported to be perturbed by established hallmarks of cancer (11). Golgi morphology in certain cancer cell types is disorganized/dispersed (12). Although in some cancer cell types, Golgi is reported to maintain cisternal organization but lack the typical ribbon organization. Also, Golgi stacks in some cancer cell types are completely disassembled or fragmented (13). Golgi organization is ‘normal’ and compact in many cancer cells, as seen in non-transformed cells (13). It is unclear if and how altered Golgi organization in cancer cells contributes to their tumorigenic characteristics or is just a consequence of cell transformation. If changes in ECM properties like stiffness could alter the organization and function of Golgi, this could contribute to anchorage independence and altered mechanosensing characteristics of cancer cells. Only one study so far has revealed links between cellular metabolism and mechanotransduction through the Golgi. It identifies neutral lipid and cholesterol synthesis as a universal response to lowered actomyosin contractility on soft matrices (14). The authors observed a reduction in extracellular mechanical forces resonates with Golgi rheology, leading to Lipin-1 inactivation affecting diacylglycerol levels at the Golgi (14). This led to reduced recruitment of Arf1 on Golgi membranes and eventual activation of transcription factors SREBP1/2 (14).

Breast cancer cell lines MCF7 and MDAMB231 are widely studied cancer models with striking phenotypic and genotypic differences in their hormone-dependency metabolic and epithelial-mesenchymal status. Previous studies have shown that anchorage-independent MCF7 and MDAMB231 cells show contrasting Golgi phenotypes in an adherent state (15). MDAMB231 cells have a predominant compact/organized Golgi, whereas MCF7 cells have a predominantly dispersed/disorganized Golgi (15). Our work has established the differential adhesion-dependent regulation of Golgi organization in these cells.

As a first step to understanding the differential regulation of Golgi organization in these cells, our studies have evaluated the differential expression of genes in the CCLE database that could localise at and regulate Golgi organization /function. This revealed differential expressed AXL in MDAMB231 vs MCF7 cells as a putative regulator of differential Golgi organization in these cells (Manuscript in bioRxiv). Our studies indicate that AXL with high expression in MDAMB231 relative to MCF7 cells localizes at the Golgi to regulate adhesion-dependent Golgi organization (Manuscript in bioRxiv). AXL is a plasma membrane-associated receptor tyrosine kinase of the TAM family (16). AXL is a known oncogene that enhances cancer cell proliferation, survival, migration, and invasion, thus promoting overall cancer progression (17). AXL overexpression and activation have also been reported in triple-negative breast cancers (18). A study by Yang et al. also showed AXL to be involved in matrix-rigidity sensing by directly controlling the local mechanosensory contractions in focal adhesions without the involvement of its ligand Gas6 (19). Few studies have explored a possible crosstalk between AXL and the Golgi apparatus. A fairly recent study has shown AXL could localize at the Golgi apparatus in highly migratory and polarized triple-negative breast cancer cells (Hs578t and MDAMB231), its targeting using a specific inhibitor, R428, affecting directed cell migration (20). A genome-wide RNAi screen study for identifying regulators of Golgi organization by targeting the kinome in HeLa cells also revealed AXL to be a possible regulator of Golgi organization (21). The physical association of AXL with Golgi-associated GTPase -Arf1 in MDAMB231 cells has also been reported. Together, these studies hint at a possible role of AXL and Arf1 in regulating Golgi organization in breast cancer cells (22). This study explored the potential role and regulation of AXL and its crosstalk with Arf1 in matrix stiffness-dependent regulation of Golgi organization in breast cancer cells.

## MATERIALS AND METHODS

### Antibodies and Reagents

#### For Western blotting and Immunostaining

Primary antibodies were diluted in the 5% BSA in 1X TBST and 1X PBS and used for western blotting and immunostaining respectively. pAkt S473 Cat. No. #4060S (1:2000), Akt Cat. No. # 9272S (1:1000). AXL Clone C89E7 Cat. No. #8661S (1:2000) and pAXL D12B2 (Y702) # Cat. No. #5724S (1:1000) was purchased from Cell Signaling. GAPDH Cat. No. #9545 (1:5000) was purchased from Sigma. GM130 Clone 35, Cat. No. # 612008 (1:100) was purchased from BD Transduction. Arf1 Clone 1D9, Cat. No. # ab2806 (1:500) was purchased from Abcam. βeta-tubulin Clone E7, Cat. No. #AB_2315513 (1:1000) was purchased from Developmental Studies Hybridoma Bank (DSHB). Anti-Acetylated Tubulin, Mouse #Cat. No. T7451 was purchased from Sigma. Phalloidin-488, Cat. No. #A12379 (1:500) was purchased from Invitrogen. DAPI, Cat no. #D1306 (0.05 mg/ml stock diluted to 1:50 times) was purchased from Invitrogen.

Secondary Fluorescent conjugated antibodies (Alexa488 and Alexa568) were purchased from Invitrogen Molecular Probes and used at a dilution of 1:1000. HRP-conjugated secondary antibodies were purchased from Jackson Immuno-research and used at a dilution of 1:5000 in 2.5 % milk made in 1X TBST.

RIPA buffer used for cell lysis was composed of the following chemicals: 50 mM Tris/HCl pH 7.5 + 0.5% sodium deoxycholate + 0.05% SDS + 150 mM EDTA pH 8.0 + 1% NP40 + 150 mM NaCl. Before lysis using the buffer, the following components were freshly added: PIC, 1 mM Sodium fluoride, and 1 mM Sodium orthovanadate.

#### For 2D polyacrylamide hydrogels

Acrylamide (HiMedia, Chennai, India, MB068), Bisacrylamide (HiMedia, MB005), APS, TEMED (Sigma, T9281), MES buffer (Sigma, M8250), NHS (Sigma, 130672), EDC (Sigma, E1769), Toluene (Qualigens, Los Angeles, CA, USA, 32507), Silane (Sigma, 440159),

Fluoromount-G (Southern Biotech, Birmingham, AL, USA, 0100-01), and Collagen (Gibco, Thermo Fisher Scientific, Waltham, MA, USA, Collagen I, Rat Tail, A1048301) were used.

#### For AXL inhibition studies

AXL kinase inhibitor-R428 (Bemcentinib and BGB324) Cat. No # 21523 was purchased from Cayman Chemical (USA).

#### Plasmids and Oligos

GalTase-RFP and Mannosidase-GFP constructs were obtained from Dr. Jennifer Lippincott Schwartz (HHMI). ABD-GFP construct was obtained from Dr. Satyajit Mayor (NCBS Bangalore), India. peGFP-N1-AXL plasmid construct was obtained from Dr. Stéphane Bodin (CRMB CNRS). pVSVG, pGagPol, AXL-pBABE puro, pBABE puro, and pmIG plasmid constructs were purchased from Addgene.

siRNA against human AXL sequence (siAXL) was designed and purchased from Merck (Sigma). siRNA sequence used were as described earlier (17, 23)

*siAXL*

*Forward* – GAAAGAAGGAGACCCGTTA

*Reverse* – TAACGGGTCTCCTTCTTTC

#### Cell culture and transfections

MDAMB231 was obtained from ECACC. MCF7 cell line was obtained from Prof. Sanjay Gupta at ACTREC, Navi Mumbai, India. All cell lines were cultured using Gibco DMEM from Thermo Fisher Scientific, adding 10% Penstrep and 5% FBS. 0.05% Trypsin was used to detach cells, and an excess culturing medium was used to neutralize the action of trypsin.

For transfection studies, cells were seeded in 6 cm dishes to attain a confluency of 60% and allowed to attach and spread for 5 hours. Using Gibco OptiMEM medium and transfection agents PEI (Sigma) and Lipofectamine 2000 (Thermo Fisher Scientific) (respectively for MCF7 and MDAMB231), the transfection mix was prepared and kept at room temperature for 30 minutes before adding to the cells seeded in 6cm dishes. Media in Lipofectamine 2000 transfected dishes was changed 12 hours post-transfection, and cells were used for experiments 36 hours post-transfection.

#### Preparation of 2D polyacrylamide hydrogels of varying stiffness

Polyacrylamide gels of varying stiffness were prepared by cross-linking 40% polyacrylamide and 2% bis-acrylamide. Briefly, gels were prepared between a glass slide uniformly coated with a hydrophobic layer of nail paint, and 12-mm glass coverslips were activated using toluene/silane solution (9: 1 ratio). Gel thickness was kept constant by using equal volumes for all gels. After polymerization, gels attached to coverslips were removed, washed with PBS, treated with 0.1 M NHS + 0.2 M EDC, and coated with 25 μg·mL^−1^ collagen overnight at 4 °C. Gels were treated with UV cycles in tissue culture hood and equilibrated in DMEM before plating cells. 10,000 and 5 × 10^5^ MDAMB231, MCF7 and AXL-MCF7 cells were seeded per gel for cell spreading and western blotting studies, respectively. Cells were allowed to spread for 24 hours and processed thereafter. For cell spreading studies, cells were fixed with 3.5% paraformaldehyde for 15 mins and stained with phalloidin at 4 °C overnight and followed by DAPI before mounting on glass slides. For western blotting, cells on each coverslip were lysed on ice with 80 μL of RIPA buffer + protease (1X PIC) and phosphatase (NaF and Sodium Orthovanadate) inhibitors. Samples were protein-estimated using Pierce™ BCA Protein Assay Kit, Cat no. #23225, and 30 μg total protein was used for western blotting.

#### AXL kinase inhibitor treatment

Post 24 hrs of cell seeding on 2D hydrogels/glass coverslips, R428 treatment was performed on cells for 12 hrs by adding the appropriate volume of the inhibitor in the culture medium so that the final concentration of R428 is 1 µM. R428 was reconstituted in DMSO as per the recommendation, and hence the same volume of DMSO was added to the culture media as control.

#### AXL knockdown using siRNA oligos

After 24 hrs of MDAMB231 cell seeding, two consecutive transfection shots were performed using RNAi max for the oligo: siAXL (50 pmol siRNA was used). The first shot was after 24 hrs of seeding, and the second one after 24 hrs of the first transfection shot. Then, cells were allowed to stay with media containing the transfection mix for another 48 hrs. Post 48 hrs, cells were harvested and seeded on the collagen-coated 2D hydrogels/ glass and were incubated for 24 hrs in 5% CO2. The remaining volume of cell suspension (approx. same cell number) was plated on 60 mm cell culture dishes for preparing lysates (post 24 hrs of replating) to validate the efficiency of the AXL knockdown using western blotting.

#### Immunofluorescence staining of cells for Golgi on 2D hydrogels and Glass

For IFA, cells fixed with 3.5 % PFA were incubated in a Permeabilization buffer for 15 minutes at room temperature. A permeabilization buffer was made with Triton X-100 (0.05%) diluted in 5% BSA and made in 1X PBS. Post permeabilization, two washes were given 1X PBS, followed by blocking with 5% BSA at room temperature for 1 hour. Three washes were given after blocking. Samples were incubated with Primary antibody overnight at 4^0^C. Three washes were given post-incubation with primary antibody followed by a 1-hour incubation with secondary fluorescent antibody at room temperature. Samples were given three washes and then mounted on slides using Fluoromount-G (Southern Biotech), Cat no. #0100-01. Slides were maintained at room temperature under dark conditions to dry, then moved to 4^0^C until confocal imaging.

#### Determining the Golgi distribution profile in a cell population

Cells where the Golgi was labelled for a cis Golgi marker GM130 were imaged using a confocal microscope; the structure of the Golgi was then observed and classified as organized, disorganized, or haze phenotypes. Representative cross-sectional confocal images of the organized and disorganized Golgi phenotypes were acquired. A minimum of 100 and a maximum of 200 randomly selected cells were observed in each phenotype population, and their Golgi structure was classified as organized or disorganized. The number of cells in each group was then used to calculate the distribution of organized versus disorganized Golgi (in percent) in each population for a given time point or treatment. Data from multiple experiments using these percentage values were plotted accordingly.

#### Microscopy imaging for cell spreading analysis

Cross section images for phalloidin stained cells were captured using 20X air objective on EVOS FL Auto Imaging System (Thermo Fisher, Waltham, MA, USA, AMAFD1000). Images for multiple frames were captured for analysis of at least 100 cells per independent experiment for respective stiffness. Cell spread analysis was performed using manual threshold using ImageJ.

#### Arf1 activity assay on 2D hydrogels (23 kPa and Glass)

MCF7 and AXL-MCF7 cells (3.5x10^5^) plated on the 2D PA hydrogels or glass coated with 25 ug/ml collagen-I for 24 hrs, were kept on ice and lysed using the activity assay buffer (24). For the lysis of cells, each PA hydrogel or glass-coated coverslip was incubated with 20 µl activity assay buffer for a total of 10 mins. Three such incubations of 10 mins each, with 20 µl activity assay buffer, were done for each gel or glass cover slip. The cells with lysis buffer were kept on ice during these treatments. At the end of 30 mins (3 incubations), all the lysate was collected from each well. To ensure an approximately equal amount of lysate protein is obtained from the lysis of cells on gels and coverslips, we used 5 x 23 kPa PA hydrogels and 3x Glass coverslips for lysis. This means we should get 5 x 60 µl = 300 µl of lysate from PA hydrogel and 3 x 60 µl=180 µl lysate from glass coverslips. To these cell lysates, an additional activity assay buffer was added to make up the total volume to 500 µl for both 23 kPa and glass. For the pulldown assay, 400 µl of these cell lysates were incubated with 60 µg of glutathione S-transferase (GST)-tagged Golgi localised γ-ear containing Arf-binding protein 3 (GGA3) fusion protein (GST-GGA3) bound to glutathione Sepharose beads. These were incubated at 4 °C on a roto-spinner at 9 rpm for 35 mins. Beads were finally spun down, washed three times with activity assay buffer and eluted in 20 µl of 2.5X Laemmli buffer. To 80 µl of the cell lysate collected at the start 20 µl of 5X Laemmli buffer was added, and this was used as whole cell lysate (WCL). The entire GGA3 pulldown eluate samples and 20 µl of WCL samples were resolved by 12.5% SDS-PAGE.

#### Generation of stable AXL expressing MCF7 cells

HEK293T cells were seeded at an appropriate density to generate retroviral particles for transduction. Post 24 hrs of HEK293T seeding, transfection using Lipofectamine 2000 was performed with pBABE -puro, pmIG and AXL-pBABE-puro plasmids along with pVSVG and pGagPol in respective dishes. Here, pMIG (a retroviral vector with the GFP-tag) was used as a control to check the transfection efficiency in HEK293T and transduction efficiency in MCF7 cells post-addition of the viral titre. After 48 hrs of transfection, 1.5x 10^5^ MCF7 cells were seeded to be used for transduction with viral titre. Also, 2 ml of 10% DMEM supplementation was done to the transfected dishes. Post 72 hrs of transfection, viral titre was collected and was filtered using the 0.2 µm filter. This was followed by the addition of 3 ml of the titre, 1 ml 10% DMEM and 4 µl of 10 µg/ml polybrene to prior seeded MCF7 cells (approx. 50% confluent). Post 72 hrs of viral titre addition, puromycin (final concentration-1.6 µg/ml) selection was performed on transduced MCF7 cells. Post 72 hrs of selection, cell viability was checked. The cell colonies surviving the selection process were replated on 24 well plate for further culturing and expansion of the colonies to confirm and check AXL protein expression by western blotting.

#### Image Analysis and quantitation

ImageJ FIJI was used to process images with scale bars for all imaging experiments. Golgi object count analysis for MCF7, MCF7 (AXL-GFP), and AXL-MCF7 cells was performed on cross-section binary images for the cis Golgi - GM130 marker (Alexa Fluor 568 channel). The Golgi object count was evaluated by manually thresholding the binary images and using the analyze particle command in ImageJ FIJI. Line plot analysis was performed using the ImageJ FIJI software to look at the overlap of markers in images. Huygens Professional software (SVI) was used to perform deconvolution of LSM files to perform colocalization analysis to look at the overlap of ABD-GFP and GM130 markers. Colocalization analysis for data showing ABD-GFP localisation at the Golgi for MCF7 and AXL-MCF7 cells was done using the Huygens software ’Colocalization Analyzer Advanced’ tool. Threshold for each cross-section image was set using the Costes method, and the GM130 (Alexa Fluor 568) channel was thresholded manually for further analysis to obtain the Pearson’s coefficient value using the software.

#### Statistics

All the statistical analysis was done using GraphPad Prism analysis software. Statistical analysis for western blotting using absolute data (not normalised to a condition or control) was done using the two-tailed unpaired Mann-Whitney U test. Two-tailed single sample t-test was used for western blot data normalised to the control. Statistical analysis for Golgi profiling data, comparing the percentage of cells with a specific Golgi phenotype across different experimental conditions, was done using one-way ANOVA, a multiple comparison test, with Tukey’s method for error correction. The Pearson’s coefficient analysis data for localisation of ABD-GFP at the Golgi was tested for statistical significance using the two-tailed unpaired Mann-Whitney U test.

## RESULTS AND DISCUSSION

### Matrix stiffness-dependent regulation of differential Golgi organization and cell spreading in MDAMB231 and MCF7

Studies from the lab have established that cell-matrix adhesion is a vital regulator of Golgi organization in ‘normal’ anchorage-dependent cells like mouse embryonic fibroblasts (10). This led us to ask how adhesion signalling affects Golgi organization in anchorage-independent cancer cells like breast cancer cell lines – MDAMB231 and MCF7, which have inherently differential Golgi phenotypes when adherent. Adherent MDAMB231 cells have a predominantly organized Golgi phenotype **(Fig 1A)**, while MCF7 cells have a predominantly disorganized Golgi phenotype **(Fig 1B)** detected using cis-medial (ManII-GFP) and trans-Golgi specific (GalTase-RFP) marker. When MDAMB231 cells held in suspension both ManII-GFP and GalTase-RFP labelled Golgi are seen to disorganise on loss of adhesion **(Fig 1A)**. MCF7 cells retain their disorganized Golgi phenotype upon loss of adhesion for both Golgi markers **(Fig 1B)**. This reveals that along with differences in their Golgi organization when these cells are adhered to the matrix, in MDAMB231, regulation of the Golgi is anchorage-dependent, while in MCF7, this regulation is anchorage-independent. Despite these differences, both the cell lines grow anchorage independently and can support tumor formation (25). This could, in part, be mediated by their ability to differentially sense and respond to mechanical cues from the matrix.

**Figure 1:**
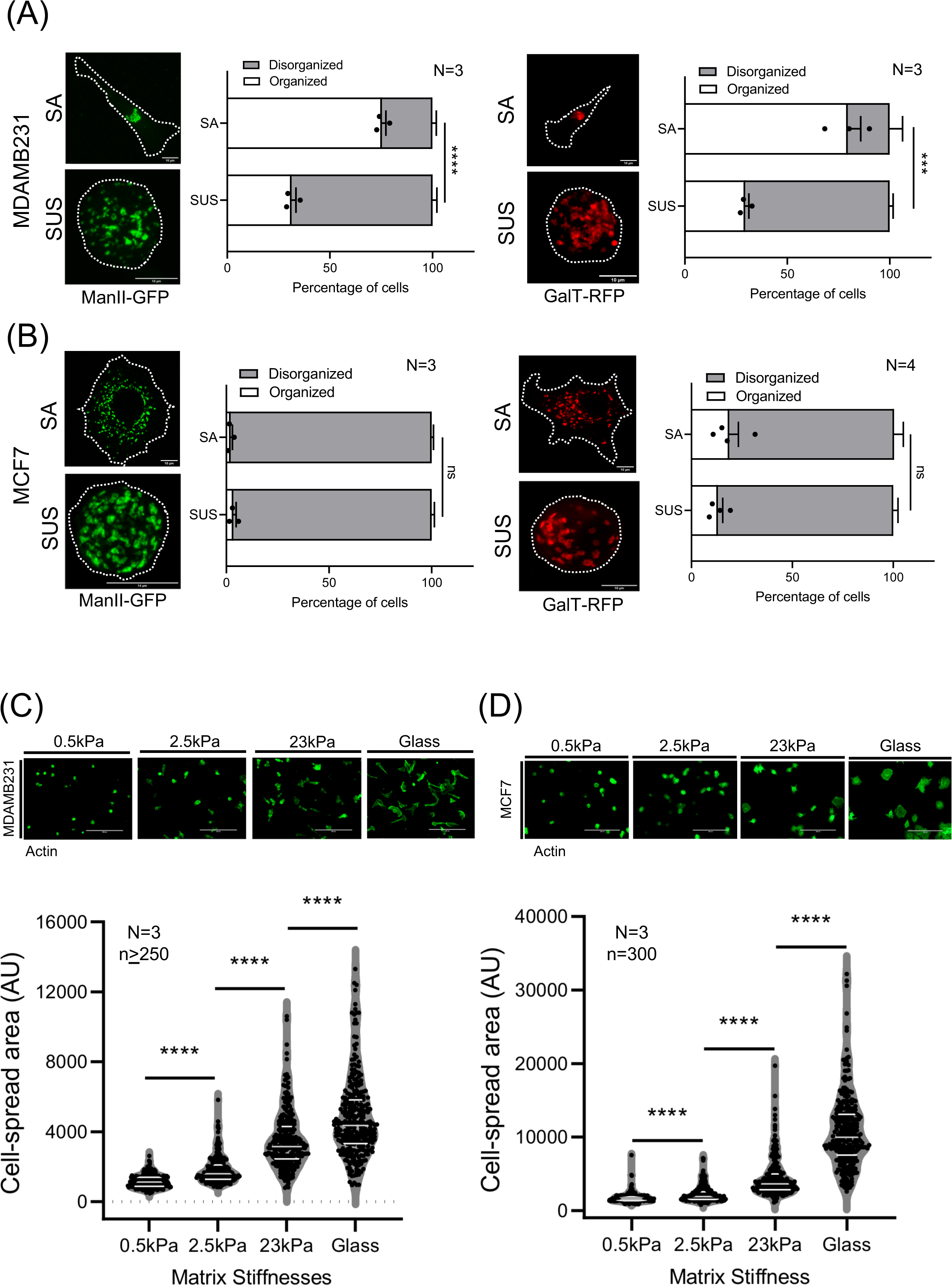
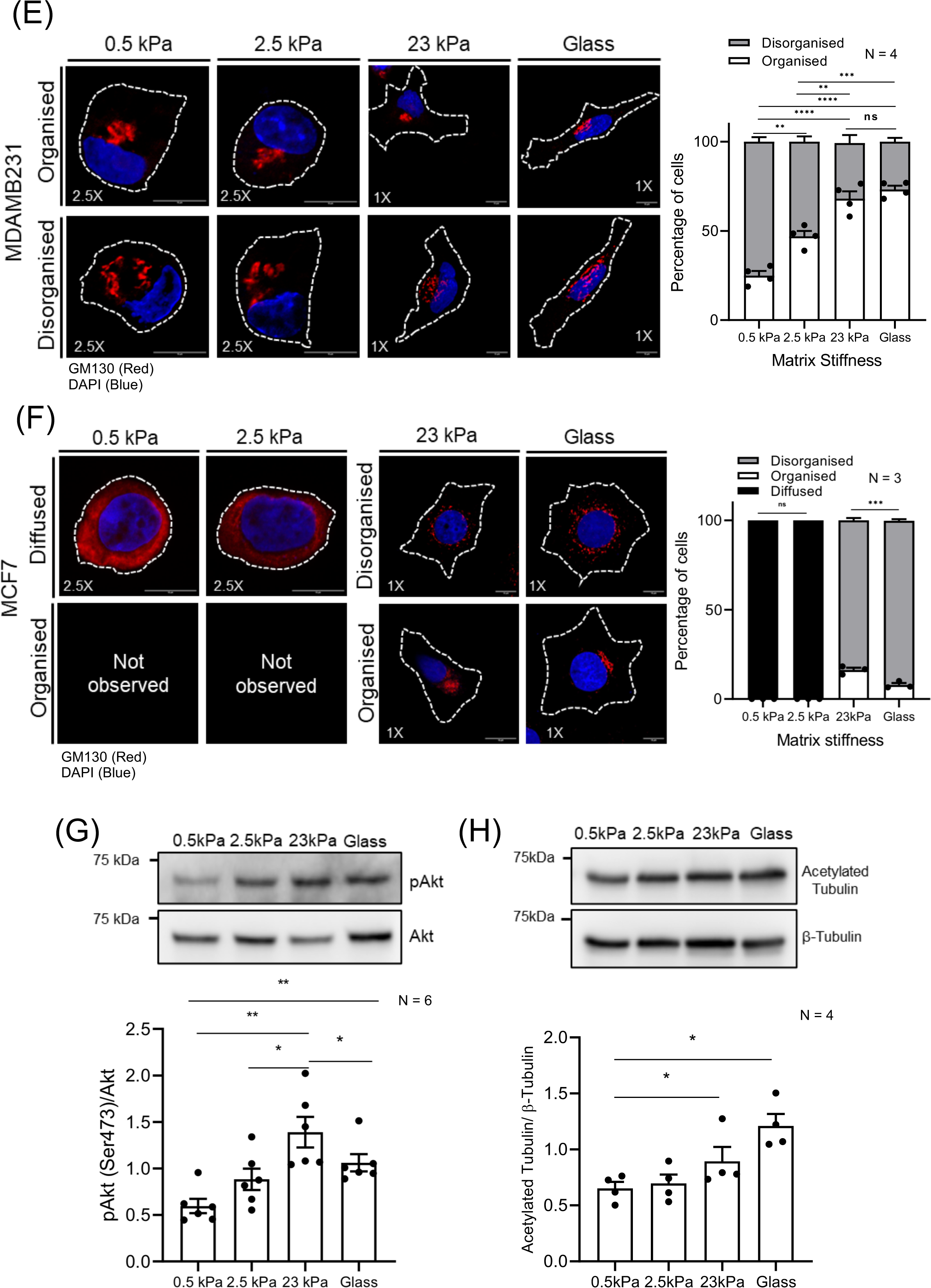
Cell-matrix adhesion-dependent matrix stiffness sensing differentially regulates Golgi organization and cell spreading in MDAMB231 & MCF7 cells. **(A, B)** Stable adherent (SA) and Suspended (SUS) MDAMB231 and MCF7 cells transfected with cis-medial Golgi (ManII-GFP) and trans Golgi (GalTase-RFP) marker and representative cross-sectional images sown. Percentage distribution profile (≥200 cells) of stable adherent (SA) and suspended (SUS) MDAMB231 and MCF7 cells with organized (white) and disorganized (Gray) Golgi. Graphs for percentage distribution profile represents mean ± SEM from three independent experiments **(C, D)** Representative cross-section images of phalloidin stained MDAMB231 and MCF7 cells adherent on collagen coated 2D polyacrylamide gels of varying stiffness (0.5kPa, 2.5kPa, 23kPa) and glass. The graph represents the median ± quartile cell spread area (represented in arbitrary units - AU) for ≥ 250 cells (n) from three independent experiment (N). **(E, F)** MDAMB231 and MCF7 cells immunostained for cis-Golgi marker GM130 (red) and nucleus labelled with DAPI (blue) on collagen coated 2D polyacrylamide gels of varying stiffness (0.5kPa, 2.5kPa, 23kPa) and glass. Representative cross-section images for cells with organised and disorganised Golgi at each stiffness are shown. Percentage distribution profile of MDAMB231 and MCF7 (n≥300) cells with Golgi that is organized (white), disorganized (Gray) and diffused with occasional puncta (black) are shown. The graphs represent mean ± SEM from four **(E)** and three **(F)** independent experiments. **(G)** Representative blots show Serine 473-phosphorylated Akt (pAkt), Akt levels and **(H)** acetylated tubulin and β-tubulin levels in MDAMB231 cells on collagen coated 2D polyacrylamide gels of varying stiffness (0.5, 2.5, 23 kPa) and glass. Graph represents the ratio of densitometric band intensities (pAkt -Ser473/Akt and Acetylated tubulin/ β-tubulin) as mean ± SEM from six (G) and four (H) independent experiments. Statistical analysis was done using the one-way ANOVA multiple comparisons test with Tukey’s method for error correction for the distribution profiles and Mann–Whitney U test for the cell spread area and western blot analysis. (*p≤0.05, **p≤ 0.01, ***p ≤ 0.001, ****p≤ 0.0001, ns=non-significant).

Studies have indeed reported that MDAMB231 and MCF7 cells differentially respond to adhesion and mechanosensory cues to regulate their migration (26), drug uptake (27) and more. To test if and how matrix stiffness-dependent mechanosensing could impact Golgi organization in these breast cancer cell lines, we first evaluated adhesion-dependent cell spreading of MDAMB231 and MCF7 cells seeded on collagen coated (25 µg/ml collagen) 2D polyacrylamide gels (0.5 kPa, 2.5 kPa, 23 kPa) and glass. Both the cell lines showed a stiffness-dependent increase in cell-spread area **(Fig 1C & D)**, confirming their mechano-responsiveness to matrix stiffness. MCF7 cells spread more than MDAMB231, especially at higher stiffness **(Fig 1C)**. This is evident when cell spread areas were plotted together and compared **(Supp Fig 1A)**.

Golgi organization profile in MDAMB231 cells (detected using cis-Golgi GM130 marker) was seen to be stiffness dependent, predominantly disorganized on softer matrices (0.5 and 2.5 kPa) and predominantly organized on stiffer matrices (23 kPa and glass) **(Fig 1E)**. These cells further show a distinct stiffness-dependent increase in Akt activation (known to be regulated downstream of integrins) that interestingly drops in cells on glass **(Fig 1G)**. The Golgi apparatus is known to be a hub for non-centrosome microtubules to nucleate (28), which, when the Golgi organization is perturbed, could be affected. In MDAMB231 cells, a stiffness-dependent increase in microtubule acetylation (and hence stability)(29) was observed **(Fig 1H)** that could possibly reflect the impact of Golgi organization being restored at higher stiffness. In MCF7 cells, the GM130 stained Golgi phenotype at lower stiffness (0.5 kPa and 2.5 kPa) is diffused throughout the cytosol with occasional punctas **(Fig 1E)**. On stiffer matrices (23 kPa and glass), the GM130-stained Golgi is predominantly disorganized, as confirmed by the distribution profile **(Fig 1E)**. In MCF7 (and MDAMB231 cells), ManII-GFP (Cis/medial Golgi marker) and GalTase-RFP (Trans Golgi marker) transfection efficiency was low, making it difficult to have enough transfected cells on 2D PA gels to evaluate the Golgi phenotype, particularly on softer matrices. Interestingly, MCF7 and MDAMB231 cells with a primary disorganised and organised Golgi phenotype respectively, have a defined secondary population with the opposite Golgi phenotype. The cause of this inherent heterogeneity and its implications for the mechano-responsiveness of these cell populations remain to be explored. Knowing what mediates matrix stiffness-dependent regulation of Golgi organization would be the first step towards this.

### Differentially expressed AXL regulates stiffness-dependent Golgi organization and cell spreading in MDAMB231 cells

AXL, a receptor tyrosine kinase known to be involved in cellular rigidity sensing (19), is differentially expressed in MDAMB231 (high expression) and MCF7 cells (no expression) **(Supp Fig 2A)**. Along with its plasma membrane localization, AXL is reported to be at the Golgi in migrating Hs578t and MDAMB231 cells (20), making it an interesting candidate for study. Using a known selective small molecule inhibitor of AXL, R428 (30), cells were allowed to adhere and adapt to gels of increasing matrix stiffness with or without the inhibitor. R428 distinctly disrupts the matrix stiffness-dependent regulation of Golgi organization in MDAMB231 cells **(Fig 2A)**. Across stiffnesses, the Golgi was disorganised in the inhibitor-treated cells **(Fig 2A).** This creates a Golgi phenotype very similar to what is observed in R428-treated MDAMB231 cells at 23 kPa and glass. R428 treatment of cells at 23 kPa and Glass showed a drop Akt activation (known to be targeted downstream of AXL and regulated by matrix stiffness) (31) confirming its effect on cells **(Supp. Fig 2B)**. R428-mediated disruption of stiffness-dependent Golgi organization in MDAMB231 also affects stiffness-dependent cell spreading. R428 treated cells spread better at all stiffnesses **(Fig 2B)**, which is evident when compared together **(Supp Fig 2C)**. This increase in spreading is seen to be prominent at higher stiffnesses (23 kPa and glass) **(Supp Fig 2C)**.

**Figure 2:**
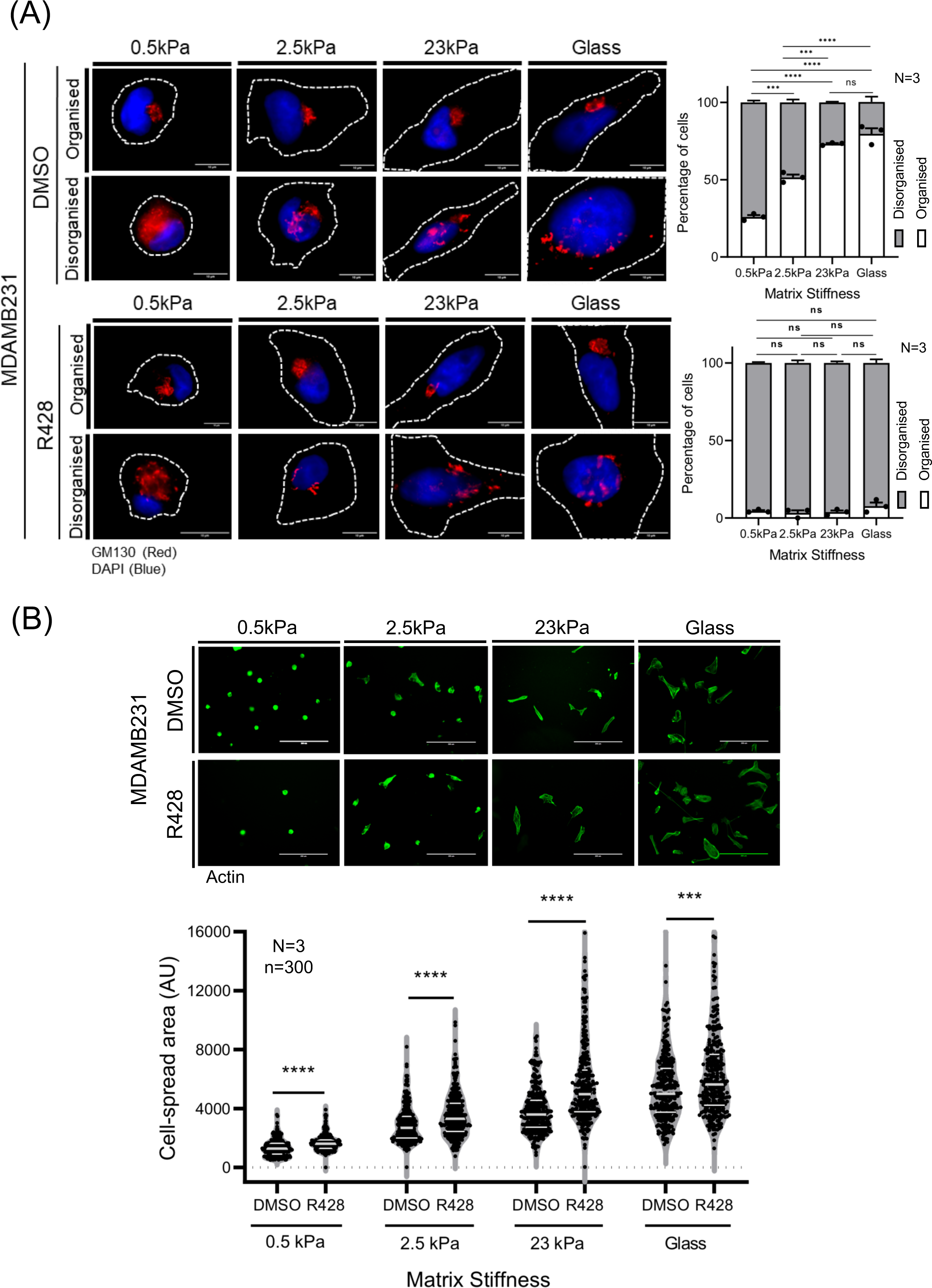
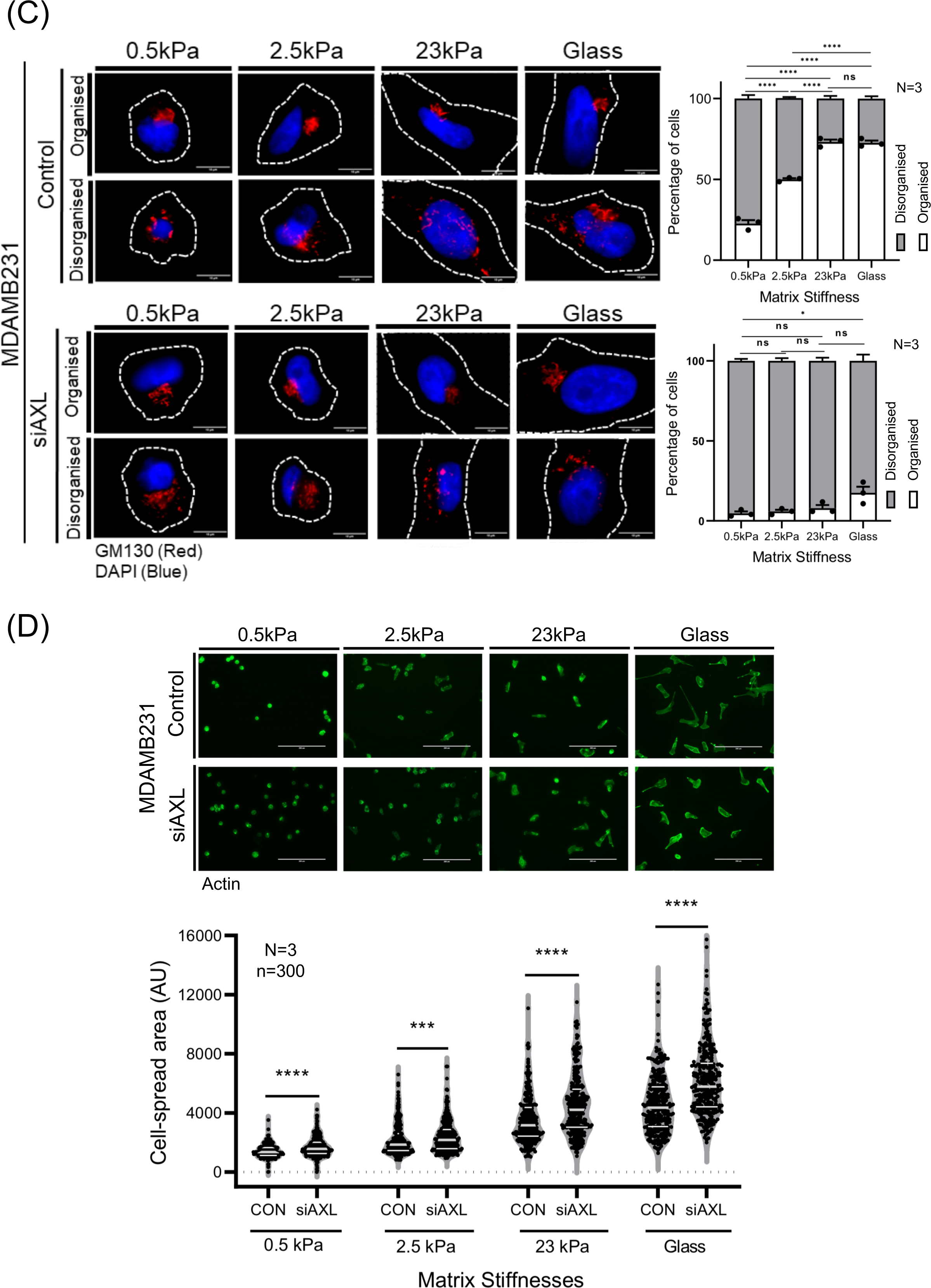
AXL activation and expression regulates stiffness-dependent Golgi organization and cell spreading in MDAMB231 cells. **(A)** Representative cross section images of DMSO and R428 treated MDAMB231 cells immunostained for cis-Golgi marker GM130 (red) and nucleus labelled with DAPI (blue) on collagen coated 2D polyacrylamide gels of varying stiffness (0.5kPa, 2.5kPa, 23kPa) and glass. Graph represents percentage distribution profile of (n≥300) cells with organized (white) and disorganized (Gray) Golgi as mean ± SEM from three independent experiments (N) **(B)** Representative cross-section images of phalloidin stained DMSO and R428 treated MDAMB231 cells adherent on collagen coated 2D polyacrylamide gels of varying stiffness (0.5kPa, 2.5kPa, 23kPa) and glass. The graph represents the median ± quartile cell spread area (represented in arbitrary units-AU) for 300 cells (n) from three independent experiment (N). **(C)** Representative cross section images of control and AXL knockdown (siAXL) MDAMB231 cells immunostained for cis Golgi marker-GM130 (red) and nucleus labelled with DAPI (blue) on collagen coated 2D polyacrylamide gels of varying stiffness (0.5kPa, 2.5kPa, 23kPa) and glass. Graph represents percentage distribution of (n≥300) cells with organized (white) and disorganized (Gray) Golgi shown as mean ± SEM from three independent experiments (N) **(D)** Representative cross-section images of phalloidin stained control and AXL knockdown (siAXL) MDAMB231 cells adherent on collagen coated 2D polyacrylamide gels of varying stiffness (0.5kPa, 2.5kPa, 23kPa) and glass. The graph represents the median ± quartile cell spread area (represented in arbitrary units-AU) for 300 cells (n) from three independent experiment (N). Statistical analysis was done using the one-way ANOVA multiple comparisons test with Tukey’s method and Mann–Whitney U test for the distribution profiles and cell spread area analysis respectively. (*p≤0.05, **p ≤ 0.01, ***p ≤ 0.001, ****p ≤ 0.0001, ns=non-significant).

We further asked if the siRNA-mediated knockdown of AXL also affects Golgi organization and cell spreading, such as in the R428 treatment. Using known tested AXL targeting siRNA sequence (17, 23), we confirmed AXL knockdown **(Supp Fig 2C)**. This, like the R428 treatment, causes the stiffness-dependent Golgi organization to be disrupted across all stiffnesses **(Fig 2C)**. AXL knockdown also causes cells to spread better across stiffness **(Fig 2D)**, which is evident when compared together **(Supp Fig 2E)**. This increase in spreading is seen to be prominent at higher stiffnesses (23 kPa and glass) **(Supp Fig 2E)** and could be mediated in part by the change in Golgi organization observed.

Together, these studies suggest expression and activation of AXL are both vital for regulating stiffness-dependent Golgi organization, which could further contribute to cell spreading in MDAMB231 cells. Earlier studies have shown Golgi-mediated trafficking and processing to be vital for cell spreading (32). In response to changing matrix stiffness, this regulation could be more finely regulated to affect cell function. With the targeting of AXL in MDAMB231, showing an MCF7-like phenotype in cells (which lack AXL), MCF7 cells could provide an ideal setting to establish the mechanosensitive regulation of Golgi organization by AXL.

### Does expression of AXL regulate Golgi organization in MCF7 cells?

All our evaluations so far on stiffness-dependent regulation of Golgi organization looked at MDAMB231 cells with high AXL expression. We know MCF7 cells inherently have very low to no AXL protein expression and a disorganised Golgi. This makes them particularly suited to test the role of AXL in regulating Golgi organization as well as cell spreading by restoring AXL expression. Transient expression of pEGFP-N1-AXL in MCF7 cells caused a reorganization of Golgi as seen by the percentage distribution profile (AXL transfected vs untransfected cells), accompanied by a significant reduction in GM130 labelled Golgi object numbers in AXL expressing cells **(Fig 3A)**. This is reflected in a significant change in the percentage distribution profile of the organized Golgi phenotype in transfected vs un-transfected cells **(Fig. 3A)**. The Golgi organization phenotype in the AXL-expressing MCF7 cells, rather than being radially distributed around the nucleus (as seen in MCF7 cells), showed a more restricted perinuclear distribution with a significant reduction in Golgi object numbers **(Fig 3A)**.

**Figure 3:**
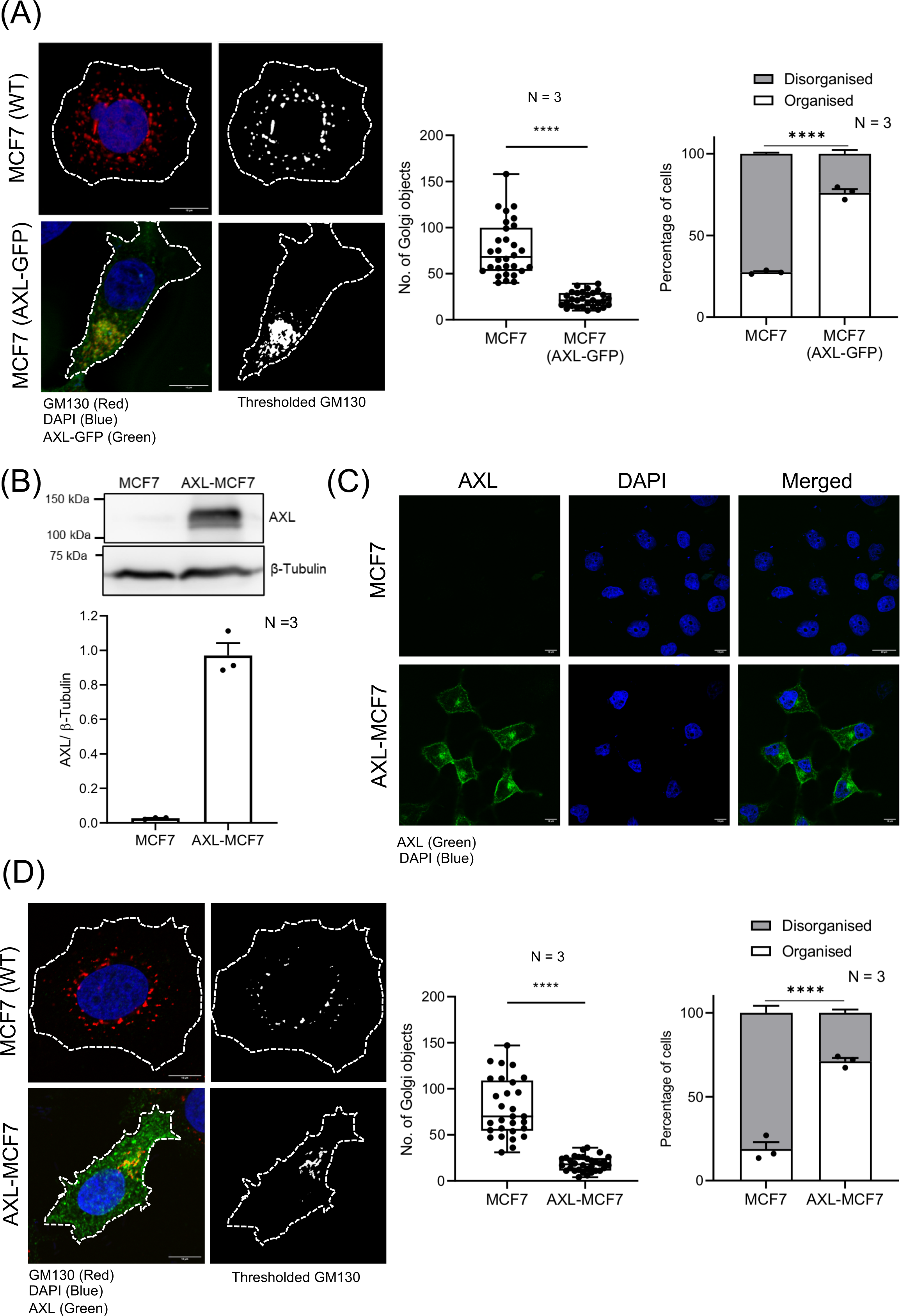
AXL expression regulates Golgi organization in adherent MCF7 cells. **(A)** Representative cross-section images for un-transfected (MCF7-WT) and AXL-GFP transfected MCF7 cells (MCF7 AXL-GFP) immunostained for cis-Golgi marker-GM130 (red) and nucleus labelled with DAPI (blue) showing the predominant Golgi organization phenotype respectively. The B/W representative cross section images show thresholded discontinuous GM130 labelled Golgi objects. The graphs for object count analysis show all data points with median ± minimum and maximum for discontinuous cis-Golgi object count per (n=30) cell and percentage distribution profile for un-transfected and AXL-GFP transfected MCF7 (n≥300) cells with organized (white) and disorganized (Gray) Golgi representing mean ± SEM, from three independent experiments (N) respectively. **(B)** Western blot detection of AXL and β-tubulin in MCF7 vs. AXL-MCF7 cells were compared. Graph represents the ratio of densitometric band intensities as mean ± SEM from three independent experiments **(C)** MCF7 and AXL expressing MCF cells (AXL-MCF7) immunostained for AXL (green) and nucleus labelled with DAPI (blue). **(D)** Representative cross-section images of similarly immunostained cells for AXL (green), cis-Golgi marker-GM130 (red) and nucleus labelled with DAPI (blue) showing the predominant Golgi organization phenotype respectively. The B/W representative cross section images show thresholded discontinuous GM130 labelled Golgi objects. The graphs for object count analysis show all data points with median ± minimum and maximum for discontinuous cis-Golgi object count per (n=30) cell and percentage distribution profile for MCF7 and AXL-MCF7 (n≥300) cells with organized (white) and disorganized (Gray) Golgi representing mean ± SEM, from three independent experiments (N) respectively. Statistical analysis was done using the one-way ANOVA multiple comparisons test with Tukey’s method for the distribution profiles and Mann–Whitney U test and object count analysis. (*p≤0.05, **p ≤ 0.01, ***p≤ 0.001, ****p ≤ 0.0001, ns=non-significant).

Stable AXL reconstitution of MCF7 cells was done using retroviral transduction. AXL expression and localization in these cells was validated by using western blotting and immunostaining **(Fig. 3B & C)**. Stable AXL expression in MCF7 cells (AXL-MCF7) restored Golgi organization accompanied by a significant reduction in Golgi object numbers relative to MCF7 cells **(Fig. 3D)**. This is reflected in a significant change in the percentage distribution profile of cells with an organized Golgi phenotype in stable AXL expressing MCF7 (AXL-MCF7) cells as compared to MCF7 cells **(Fig 3D)**. Golgi organization in stable AXL-MCF7 cells was comparable to transient AXL expression **(Fig. 3A & D)**. This was confirmed using ManII-GFP (Cis/medial) and GalTase-RFP (Trans) Golgi markers **(Supp Fig. 3B)**. Together, they confirm AXL expression to be important for the differential Golgi organization in MDAMB231 and MCF7 cells.

Next, we checked if AXL-MCF7 cells respond to changing matrix stiffness, like MDAMB231 cells (expressing AXL). In MCF7 cells, the GM130 stained Golgi phenotype at lower stiffness (0.5 kPa and 2.5 kPa) was diffused throughout the cytosol with occasional puncta observed **(Fig. 1E)**. On stiffer matrices (23 kPa and glass), the Golgi phenotype is predominantly disorganized **(Fig. 1E) (Fig 4A)**. In AXL-MCF7 cells, the Golgi phenotype is restored to being predominantly organised at 23 kPa and Glass **(Fig. 4A)**. At lower stiffnesses (0.5 and 2.5 kPa), the Golgi organization stayed diffused with occasional puncta **(Fig. 4A)**. This suggests in MCF7 cells at lower stiffness, AXL may not be the primary or sole player to regulate the Golgi organization. We next asked if AXL expression and resulting regulation of Golgi organization in AXL-MCF7 cells affects cell spreading in response to increasing stiffness. AXL-MCF7 cells spread significantly less compared to MCF7 cells across all stiffnesses **(Fig 4B & Supp. Fig 4A)**. Together, they suggest AXL expression/activation to regulate stiffness-dependent cell spreading in MCF7 **(Fig. 4B)** and MDAMB231 cells **(Supp Fig 2C & 2E)**. MCF7 studies show at lower stiffness, AXL-dependent cell spreading could be regulated independently of Golgi organization **(Fig. 4A & B)**. AXL-mediated regulation of Golgi organization in response to changing matrix stiffness could be regulated by adhesion-dependent signalling. Adhesion-dependent regulation of Arf1 is known to control Golgi organization (10). Arf1 activation is shown to be regulated by actomyosin contractility in cells

**Figure 4:**
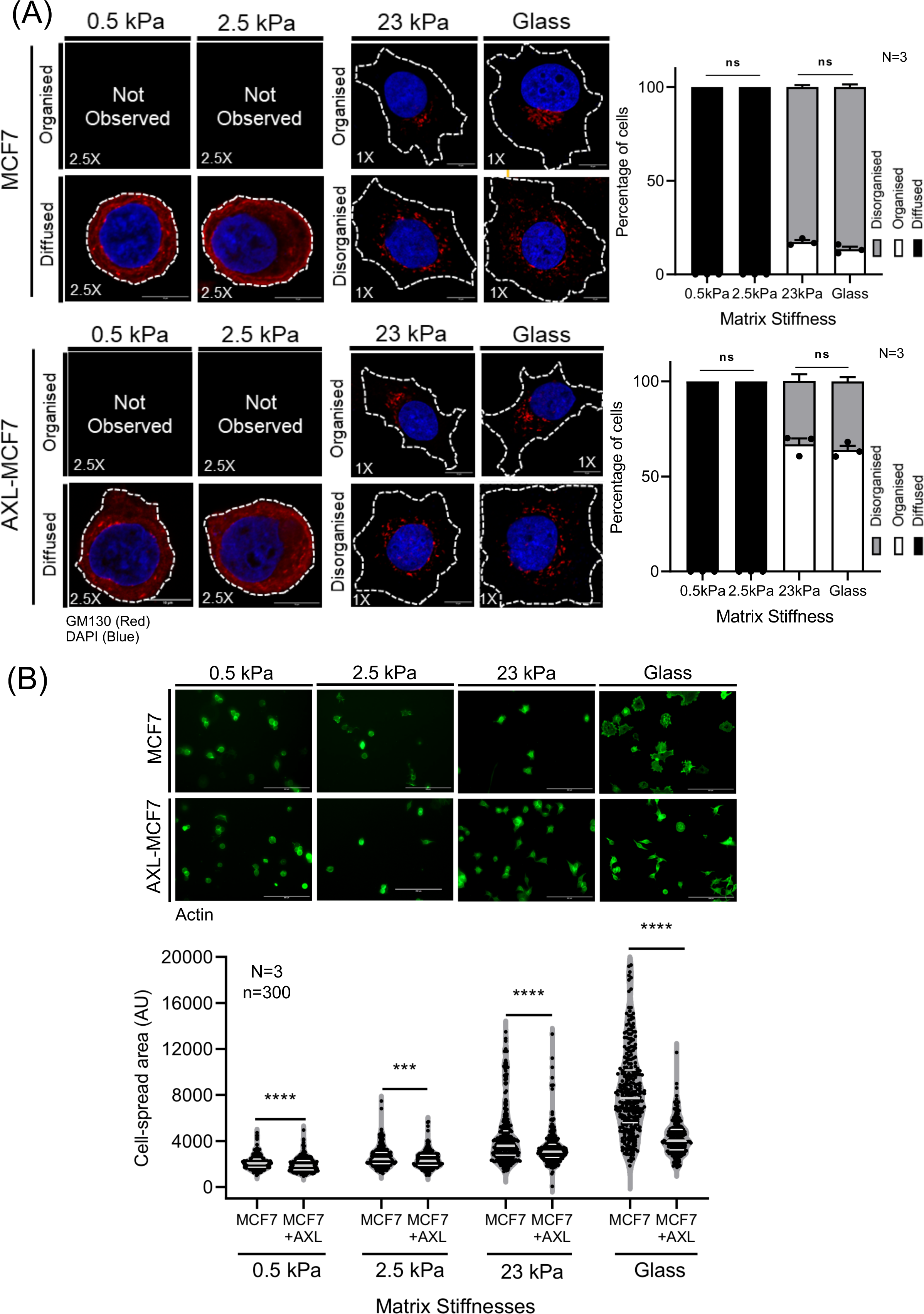
AXL expression regulates Golgi organization and cell spreading in MCF7 cells on stiffer matrices. **(A)** Representative cross section images of MCF7 and AXL expressing MCF7 cells (AXL-MCF7) immunostained for cis Golgi marker - GM130 (red) and nucleus labelled with DAPI (blue) on collagen coated 2D polyacrylamide gels of varying stiffness (0.5kPa, 2.5kPa, 23kPa) and glass. Graphs represent percentage distribution profile of (n≥300) cells with organized (white) and disorganized (Gray) Golgi as mean ± SEM from three independent experiments (N) **(B)** Representative cross-section images of phalloidin stained MCF7 and AXL-MCF7 cells adherent on collagen coated 2D polyacrylamide gels of varying stiffness (0.5kPa, 2.5kPa, 23kPa) and glass. The graph represents the median ± quartile cell spread area (represented in arbitrary units - AU) for 300 cells (n) from three independent experiment (N). Statistical analysis was done using the one-way ANOVA multiple comparisons test with Tukey’s method for the distribution profiles and Mann–Whitney U test and cell spread area analysis. (*p≤0.05, **p≤ 0.01, ***p ≤ 0.001, ****p ≤ 0.0001, ns=non-significant).

(14). ROCK and MLCK inhibition in cells are seen to cause a drop in Arf1 activation (14). This could mimic their behaviour on a softer matrix. Arf1 has also been shown to interact with AXL (22) and could possibly be regulated by it, making their crosstalk an attractive pathway to test.

### Role of AXL-Arf1 crosstalk in stiffness-dependent regulation of Golgi organization

Previous studies have shown Arf1 activation and localization at the Golgi to be vital for regulating its organization and functions (33)(34)(35). We found adhesion-dependent Arf1 activation to regulate Golgi organization and function in anchorage-dependent mouse fibroblasts (36). Active Arf1 can also bind AXL in MDAMB231 cells in a previous study (22). This makes an adhesion-AXL-Arf1-Golgi pathway of particular interest. We hence evaluated the expression of AXL and Arf1 in MDAMB231 cells across varying matrix stiffness. AXL showed a stiffness-dependent increase in expression levels **(Fig. 5A)** without a change in AXL phosphorylation (Y702) **(Supp.** Fig. 5A). Net pAXL (Y702) levels, however, increase across stiffness possibly reflecting the change in net AXL levels **(Supp. Fig. 5B)**. The effect Y702 AXL phosphorylation has on AXL activation status is variable in literature, making its interpretation challenging (30, 37, 38). Independent studies with AXL inhibitor R428 from the lab do suggest inhibitor R428 causes an increase in pY702 AXL levels in MDAMB231 and A549 cells (Manuscript in bioRxiv). Total Arf1 levels also show a simultaneous matrix stiffness-dependent increase in MDAMB231 cells **(Fig. 5B)**. In MCF7 cells lacking AXL, this matrix stiffness-dependent increase in Arf1 levels was interestingly not seen **(Fig 5C)**. Together, the stiffness-dependent regulation of AXL and Arf1 expression only further suggests their regulatory crosstalk could also be of interest in the stiffness-dependent regulation of Golgi organization.

**Figure 5:**
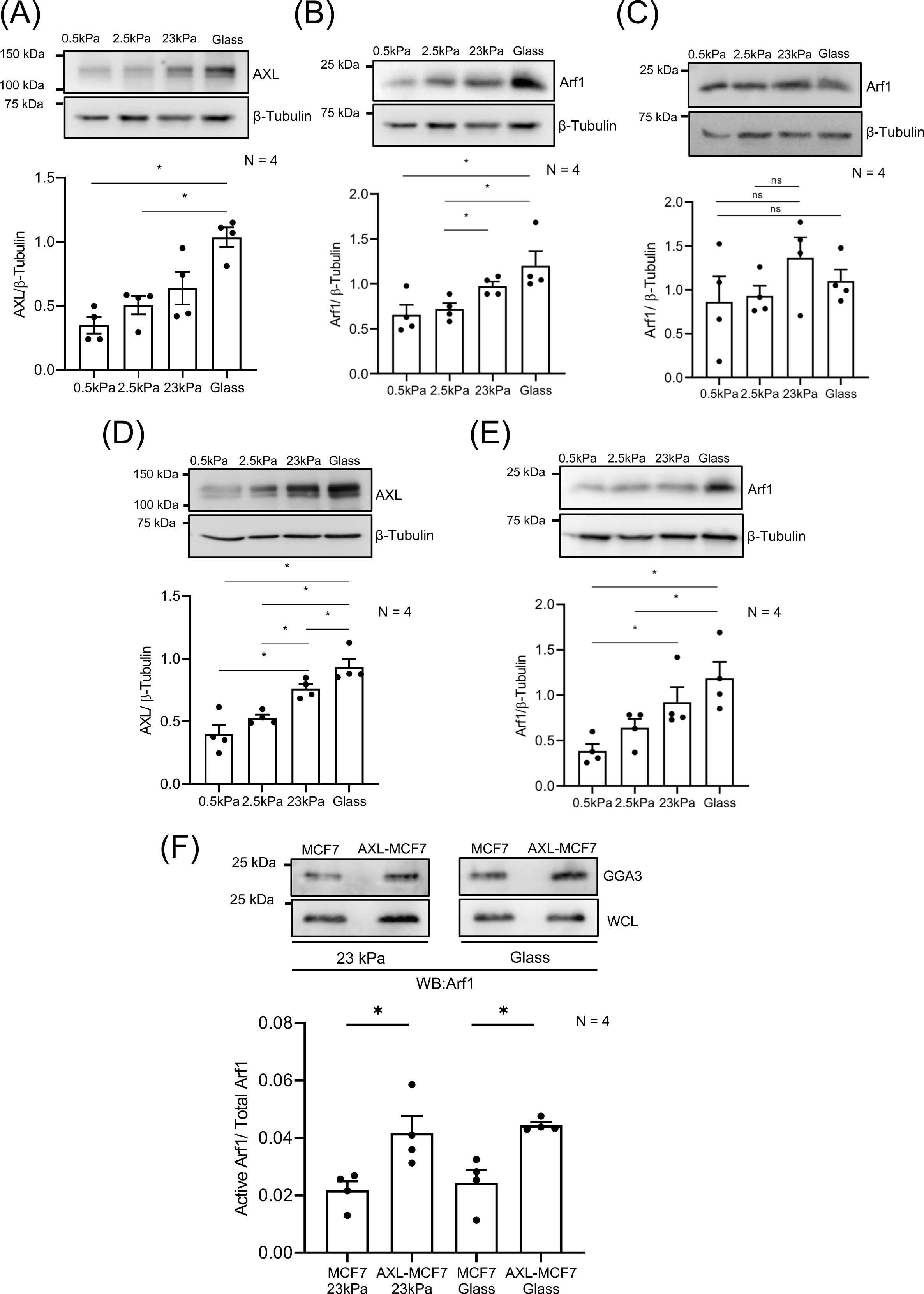
AXL-Arf1 crosstalk in regulating stiffness-dependent Golgi organization in MDAMB231 and MCF7 cells. Western blot detection of AXL levels in cell lysates from **(A)** MDAMB231 **(D)** AXL-MCF7 cells and Arf1 levels from **(B)** MDAMB231 **(C)** MCF7 and **(E)** AXL-MCF7 cells on collagen coated 2D polyacrylamide gels of varying stiffness (0.5, 2.5, 23 kPa) and glass. Graphs represent the ratio of densitometric band intensities as mean ± SEM from four independent experiments **(F)** Active Arf1 pulled down using GST-GGA3 (GGA3) and total Arf1 in whole cell lysate-WCL) from MCF7 and AXL-MCF7 cells on collagen coated 23 kPa and glass. Graph represents the ratio of densitometric band intensities as mean ± SEM from four independent experiments. Statistical analysis was done using Mann-Whitney U test. (*p≤0.05, **p ≤ 0.01, ***p ≤ 0.001, ****p ≤ 0.0001, ns=non-significant).

To test this regulatory crosstalk, we evaluated the matrix stiffness-dependent regulation of AXL-MCF7 cells. Basal Arf1 levels of MCF7 and AXL-MCF7 cells showed no difference when compared **(Supp. Fig 5C)**. AXL-MCF7 cells showed a matrix stiffness-dependent increase in AXL **(Fig. 5D)** and Arf1 levels **(Fig. 5E)** like MDAMB231 cells. This does suggest a regulatory overlap in expression between AXL and Arf1 that could further reflect in functional crosstalk vital for Golgi organization at higher stiffnesses (23 kPa and Glass). Arf1 activation status in these cells is known to be vital in regulating adhesion-dependent Golgi organization. This could partly be mediated by the stiffness-dependent increase in net Arf1 levels causing an increase in active Arf1 levels. The presence of AXL could also impact Arf1 activation in these cells. To evaluate this, we compared active Arf1 levels (using a GST-GGA3 pulldown assay) in MCF7 and AXL-MCF7 cells on 23 kPa and glass. Active Arf1 levels were significantly higher in AXL-MCF7 than in MCF7 cells on 23 kPa and glass **(Fig. 5F)**. Interestingly, active Arf1 detected using ABD-GFP in AXL-MCF7 cells on glass extensively colocalises with GM130-stained Golgi, unlike in MCF7 cells where the Golgi is disorganised **(Supp. Fig 5E)**. Together, they reveal that on higher stiffness, AXL, along with regulating Arf1 levels, also regulates Arf1 activation at Golgi to drive its organization. This makes AXL and its crosstalk with Arf1 vital mechanosensory regulators of Golgi organization in breast cancer cells.

## DISCUSSION

Among the several well-established hallmarks of breast cancers, stiffening of the tumor matrix is now widely known to promote breast tumorigenesis (39). Due to changes in the breast tumor microenvironment, cells must adapt to the altered stiffness conditions. This, in turn, supports cancer cell growth, metabolism and migration, which drives tumorigenesis (40–42). MDAMB231 and MCF7 cells differ in their mechanical properties and invasive behaviour (43) making them an attractive system for studying the role and regulation of cellular mechanosensing. This is largely thought to be mediated by their differential regulation of the actin cytoskeleton (44). A distinct difference in the Golgi organization of MDAMB231 (organized Golgi) and MCF7 (disorganized Golgi), while known, has not been explored as a regulator of their differential mechanoresponsiveness. Despite cell-matrix adhesion being known to regulate Golgi organization and function (10) only one study so far has evaluated the mechanosensitive nature of the Golgi (45). Our earlier studies have shown that matrix-dependent Golgi organization is regulated by Arf1 activation (10). Changes in extracellular mechanical forces affect Golgi rheology, leading to Lipin-1 inactivation and alterations of diacylglycerol levels at the Golgi (14). This is also mediated by Arf1-dependent regulation of transcription factors like SREBP1/2 (14). Studies have shown that elevated Arf1 levels and activation plays a key role in the hyper-proliferative behaviours of breast cancer cells (46, 47). Our independent studies evaluating the differential expression of Golgi-associated genes in MDAMB231 vs MCF7 cells identified AXL as one of the top regulatory candidates, revealing an AXL-Arf1-Golgi regulatory pathway (Manuscript in bioRxiv).

On loss of adhesion, the Golgi in MDAMB231, which becomes disorganised as compared to MCF7, where it stays disorganised, could hence reflect on changes that ensue when they respond to changing matrix stiffness. The increasing percentage of MDAMB231 cells with an organised Golgi on increasing stiffness is the first report of its kind to indicate that Golgi organization in breast cancer cells could be sensitive to matrix stiffness. The heterogeneity in Golgi organization phenotypes on each stiffness is conserved and could have implications for how the cell population behaves at that stiffness. Do cells with differing Golgi organization at the same stiffness vary in Golgi function? This remains to be tested. Loss of AXL protein or its activation (R428) were both seen to dramatically disrupt Golgi organization and its responsiveness to changing matrix stiffness. This strongly supports the role of AXL as a mechanosensitive regulator of Golgi organization in these cells.

In MDAMB231 cells, AXL expression is seen to respond to changing matrix stiffness, as is the case with Arf1. AXL, when expressed in MCF7 cells, also responds similarly, as does Arf1. The known regulatory crosstalk between YAP and AXL could mediate this (48). YAP overexpression in head and neck squamous cell carcinoma (HNSCC) promotes AXL expression (48). AXL is also seen to regulate YAP and its oncogenic function (49). The impact AXL expression has on Arf1 expression could also be mediated through its regulation of YAP in these cells (49, 50). This matrix stiffness-dependent regulation of AXL and Arf1 could have implications beyond their regulation of Golgi in breast cancer cells. Could such regulatory crosstalk exist in other cells? Or is this unique to breast cancer? This is an open question. AXL-mediated regulation of Arf1 activation is normalised for change in Arf1 expression, suggesting this to be an additional element to their crosstalk. In our independent studies, the inhibition of AXL and Arf1 are both seen to cause them to leave the Golgi in MDAMB231 cells. This suggests their regulation of each other’s activation could impact their localization and function (Manuscript in bioRxiv). The AXL high-affinity ligand; Gas6-mediated signalling regulates tumour growth, metastasis, invasion, epithelial-mesenchymal transition (EMT), angiogenesis, drug resistance, immune regulation, and stem cell maintenance (51). The implications AXL-Gas6 binding at the plasma membrane could have on its role at the Golgi could further add to our understanding of the AXL-Arf1-Golgi crosstalk. Ongoing work in the lab is working to address the same.

In MCF7 (lacking AXL) and AXL-MCF cells on softer matrices (0.5 kPa and 2.5 kPa), the Golgi phenotype is strikingly different, with a unique diffused distribution of cis-Golgi marker GM130. Knowing the localization of GM130 is closely associated with the cis-Golgi it is a strong indication that Golgi organization is dramatically altered in these cells. Challenges in imaging cells on softer gels made characterising this phenotype difficult. Knowing AXL and Arf1 expression to show a small change between 0.5 kPa and 2.5 kPa suggests that their expression is not prominently regulated at this stiffness range. This could imply that additional changes in MCF7 may regulate Golgi’s organization at a lower stiffness. This is also distinctly different from MDAMB231, where even in the absence of AXL the Golgi does not have a diffused organization at lower matrix stiffness. This suggests there could be more to the mechanosensitive regulation of Golgi in breast cancer cells, particularly in the case of MCF7 cells.

AXL expression and inhibition are further seen to regulate stiffness-dependent cell spreading of breast cancer cells. In the absence of AXL, cells with a dispersed Golgi spread more. AXL is also known to regulate focal adhesion dynamics (52), which could be responsible for the changes observed. Golgi organization in regulating trafficking could contribute to focal adhesion-dynamics. Golgi associated microtubules interact with focal adhesions to regulate cell polarization and durotaxis on substrates of increasing stiffness (53). Changes in MT-acetylation could be mediated by the regulation of Arf1 levels/activation (54) and Golgi organization (28), both regulated by changing matrix stiffness and are dependent on AXL expression/activation.

Studies have implicated changes in cell glycan signatures as a significant factor for breast cancer tumorigenesis (55). As tumors become more metastatic and invasive, specific N and O-glycan changes are observed on glycosylated proteins and lipids trafficked through the Golgi to the plasma membrane (56, 57). These changes in glycan signatures are strongly linked with alterations in Golgi organization and function (58). Studies have also hinted towards the possibility of matrix stiffness affecting the surface glycans of cells called the glycocalyx (59), which could also be affected by Golgi organization and function. Breast cancer aggressiveness is significantly dependent on the stiffness of the extracellular matrix (ECM), with stiffer matrices correlating with more aggressive tumour behaviour, promoting invasion and metastasis (39). The AXL-Arf1-Golgi pathway could hence prove vital to these changes associated with breast cancer tumorigenesis and its regulation by matrix stiffness.

## Acknowledgement.

This work is supported by a SERB CRG grant – CRG/2022/001813 to NB. AS is supported by a Prime Minister’s Research Fellowship (PMRF). We acknowledge the extensive support provided by the IISER Pune Microscopy Facility for cellular imaging. We also acknowledge the Prof. Anshuman Nag lab (IISER, Pune) for their help with and use of the spin coater. We acknowledge the help of Prachi Joshi and Radhika Malaviya for their reading of and comments on the manuscript.

## Author Contributions

The experimental work was led jointly by AS and TS. Data was recorded, organized and analyzed by TS and KS. The manuscript was written through the contributions of all authors. All authors have given approval to the final version of the manuscript. The authors declare no competing financial interest.

## SUPPLEMENTARY FIGURE LEGENDS

**Supplementary Figure 1: (A)** Cell spread area trends of MCF7 and MDAMB231 on collagen coated 2D polyacrylamide gels of varying stiffness (0.5, 2.5, 23 kPa and glass). The graph represents the trend for cell spread area plotted as median with 95% CI (error bar). Statistical analysis was done using Mann-Whitney U test. (*p≤0.05, **p ≤ 0.01, ***p ≤ 0.001, ***p ≤ 0.0001, ns=non-significant).

**Supplementary Figure 2: (A)** Representative western blot of AXL and β-tubulin levels in MDAMB231 vs MCF7 cells. **(B)** Western blot detection of Phospho-Akt (pAkt Ser473) and Akt in DMSO and R428 treated MDAMB231 cells on 23 kPa polyacrylamide gel and glass. Graph represents the ratio of densitometric band intensities (pAkt Ser473/Akt) as mean ± SEM from four independent experiments. **(C)** Cell spread area trends of DMSO and R428 treated MDAMB231 cells on collagen coated 2D polyacrylamide gels of varying stiffness (0.5, 2.5, 23 kPa) and glass. The graph represents the trend for cell spread area plotted as median and 95% CI (error bar). **(D)** Western blot detection of AXL and β-tubulin in control and AXL knockdown (siAXL) in MDAMB231 cells. Graph represents the ratio of densitometric band intensities (normalised to control) as mean ± SEM from three independent experiments. **(E)** Cell spread area trends of control and AXL knockdown (siAXL) in MDAMB231 cells on collagen coated 2D polyacrylamide gels of varying stiffness (0.5, 2.5, 23 kPa) and glass. The graph represents the trend for cell spread area plotted as median with 95% CI (error bar). Statistical analysis was done using Mann-Whitney U test for non-normalised western blot and cell spread area analysis. Single sample Wilcoxon t test was used for normalised (with respect to control) western blotting analysis. (*p≤0.05, **p ≤ 0.01, ***p ≤ 0.001, ****p ≤ 0.0001, ns=non-significant).

**Supplementary Figure 3: (A)** Brightfield representative images to show the comparative cell morphology of MCF7 and AXL-MCF7 cells **(B)** Stable adherent AXL-MCF7 cells transfected with cis-medial Golgi (ManII-GFP) and trans Golgi (GalTase-RFP) marker and immunostained for AXL. Representative cross-section images and percentage distribution profile (n≥150 cells) of adherent AXL-MCF7 cells with organized (white) and disorganized (Gray) Golgi. Graph represents data from one experiment.

**Supplementary Figure 4: (A)** Cell spread area trends of MCF7 and AXL-MCF7 cells on collagen coated 2D polyacrylamide gels of varying stiffness (0.5, 2.5, 23 kPa) and glass. The graph represents the trend for cell spread area plotted as median with 95% CI (error bar). Statistical analysis was done using Mann-Whitney U test. (*p≤0.05, **p ≤ 0.01, ***p ≤ 0.001, ****p ≤ 0.0001, ns=non-significant).

**Supplementary Figure 5:** Western blot detection of Phospho-AXL **(A)** (pAXL-Y702) to total AXL levels and **(B)** Arf1 to β-Tubulin levels in MDAMB231 cells on collagen coated 2D polyacrylamide gels of varying stiffness (0.5, 2.5, 23 kPa) and glass. Graphs represent the ratio of densitometric band intensities as mean ± SEM from four independent experiments respectively. Western blot detection of **(C)** Arf1 and β-Tubulin and **(D)** Phospho-Akt (pAkt Ser473) and Akt levels in MCF7 vs AXL-MCF7 cells. Graphs represent the ratio of densitometric band intensities as mean ± SEM from three independent experiments respectively. **(E)** Representative cross-section images and line plot analysis of adherent MCF7 and AXL-MCF7 cells expressing ABD-GFP (green) for detecting active Arf1 and immunostained for GM130 (red). Graph represents the Pearson’s correlation coefficient analysis performed on deconvoluted cross section images for colocalization of ABD-GFP (green) and GM130 (red) plotted as mean ± SEM for n=10 and 11 cells. Statistical analysis was done using Mann–Whitney U test for western blot and colocalization analysis. (*p≤0.05, **p ≤ 0.01, ***p ≤ 0.001, ****p ≤ 0.0001, ns=non-significant).

