## Supplementary figures and images for "Differential AXL expression and regulation of Arf1 controls matrix stiffness-dependent Golgi organization and function in breast cancer cells"

### Supplementary Figures

Supplementary Fig 1

(A)

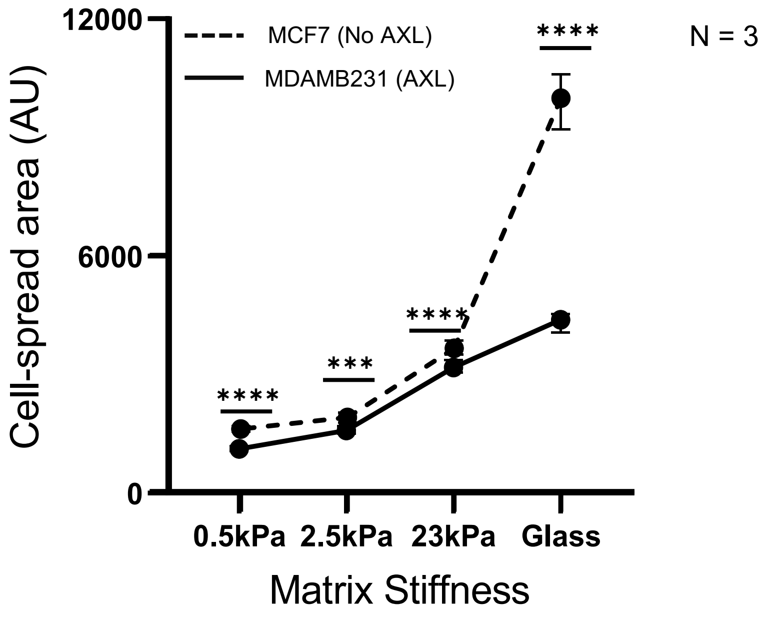

Supplementary Fig 2

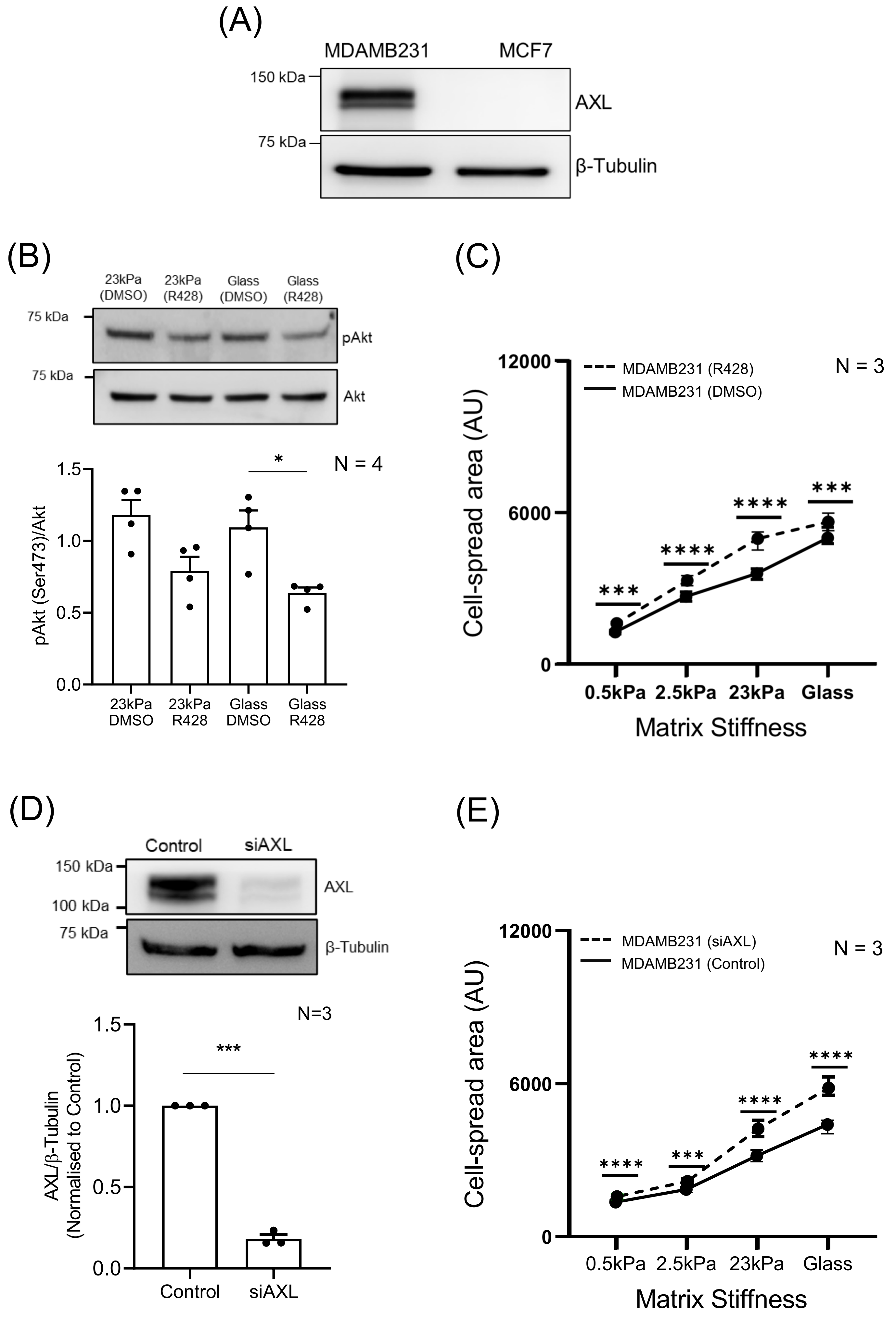

Supplementary Fig 3

(A)

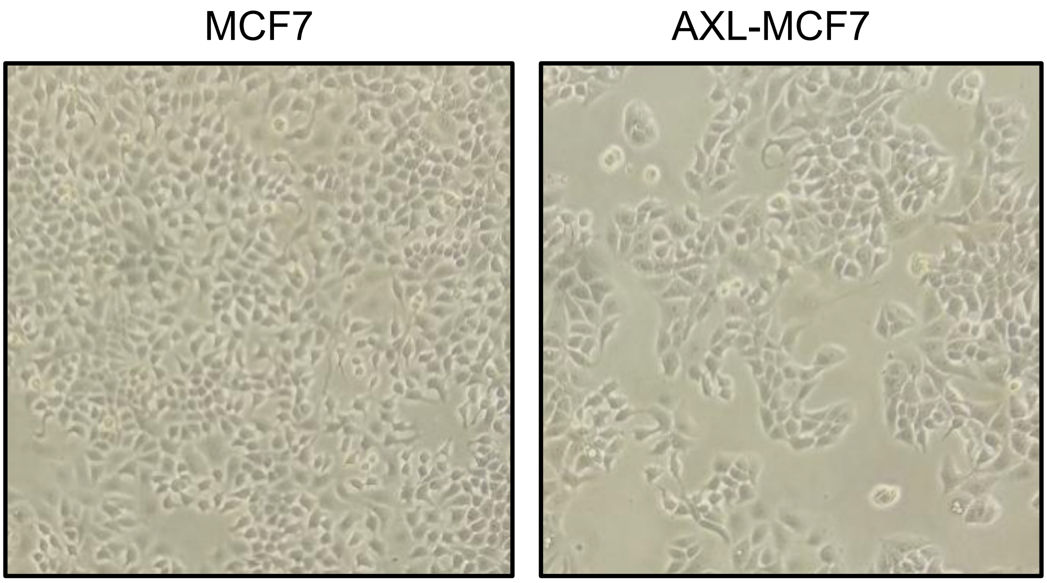


(B)

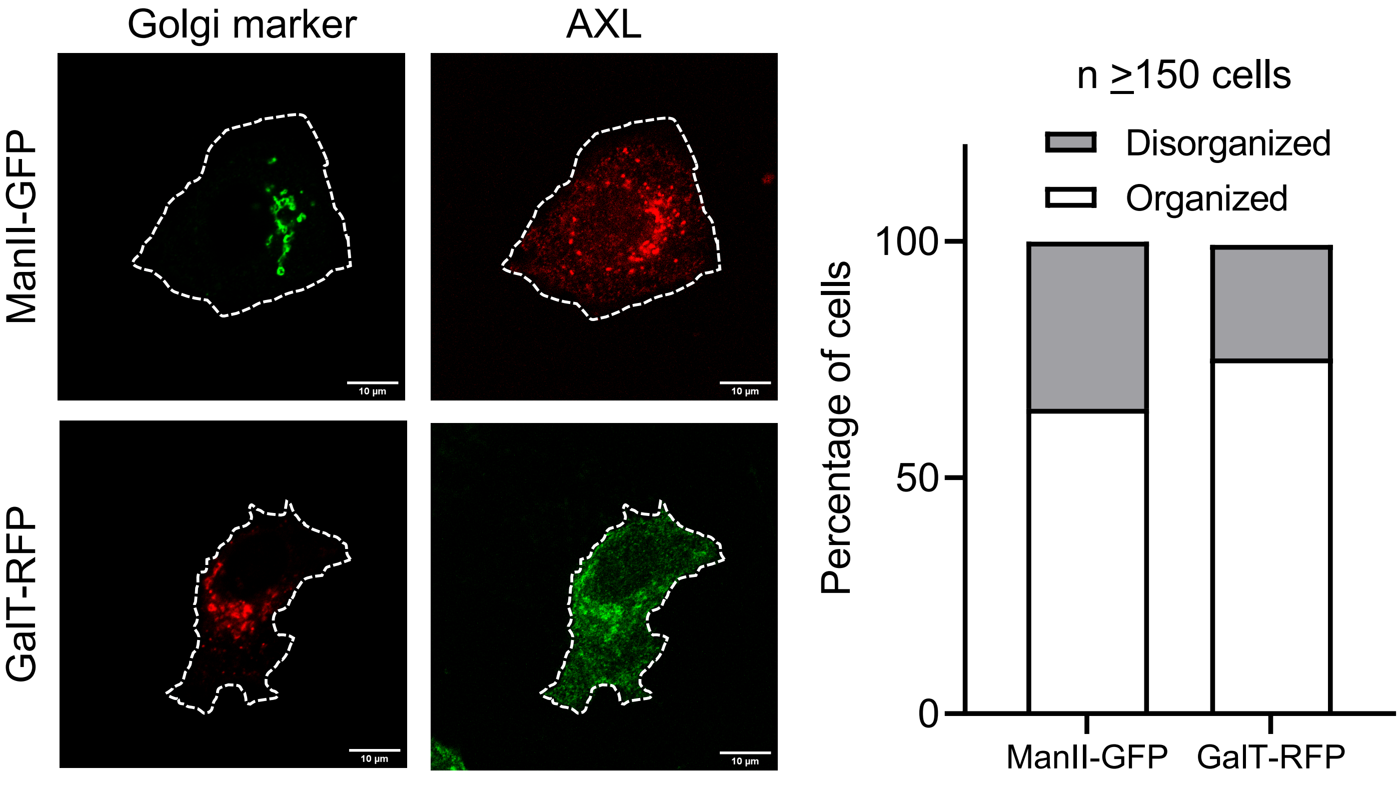

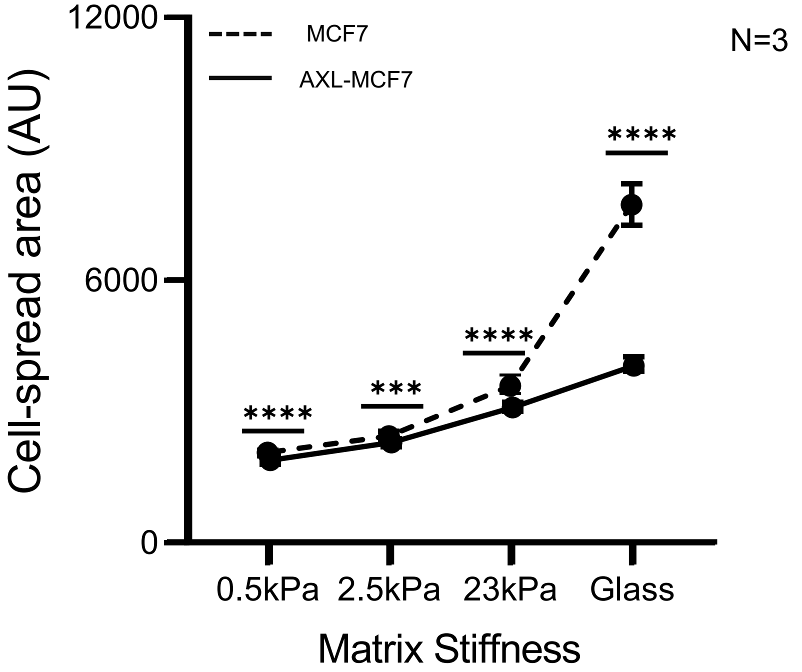

Supplementary Fig 5

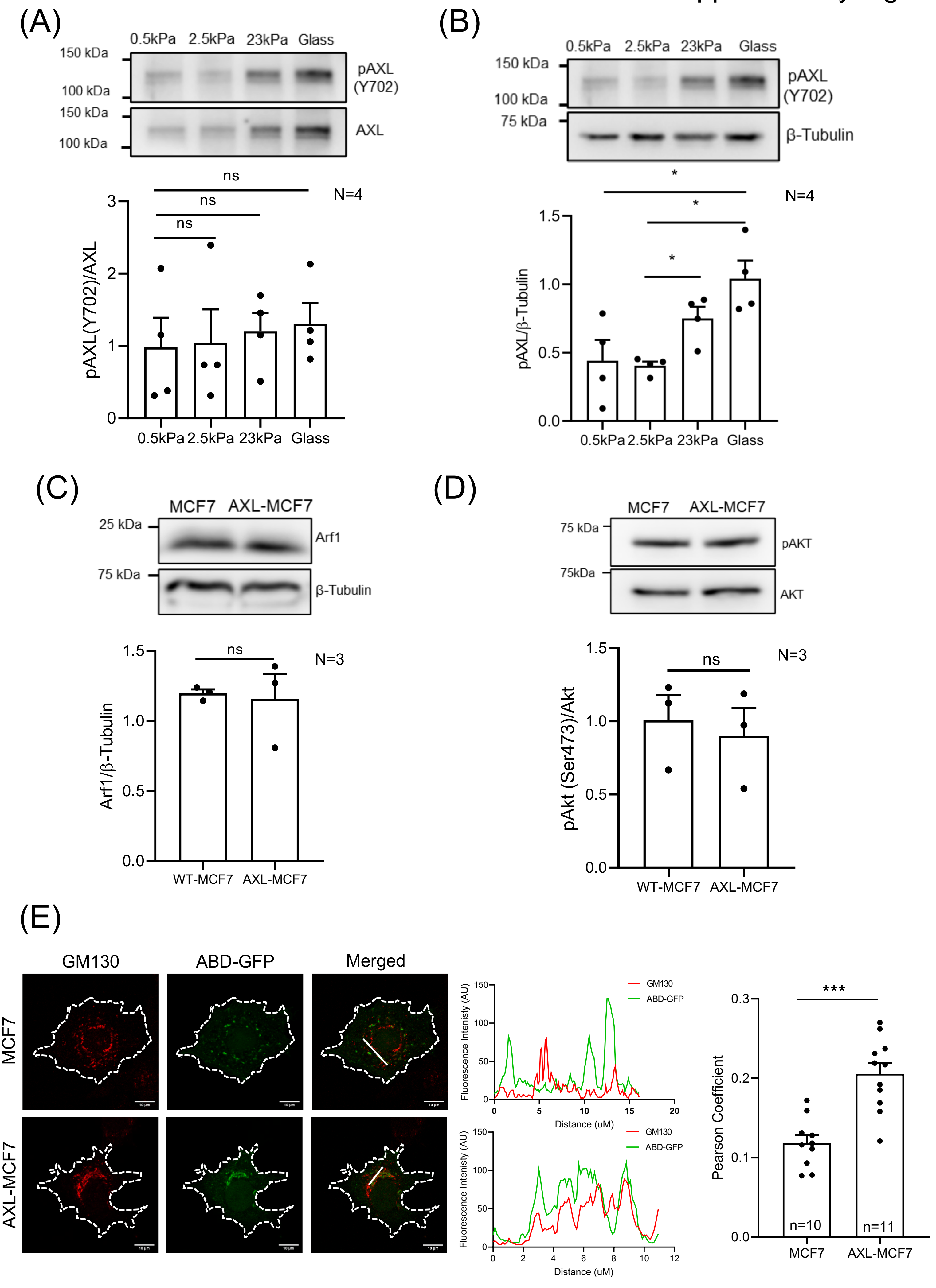
